# Application of EGCG modified EDC/NHS cross-linked extracellular matrix to promote macrophage adhesion

**DOI:** 10.1101/2020.10.03.325068

**Authors:** Chenyu Chu, Shengan Rung, Renli Yang, Yi Man, Yili Qu

## Abstract

Though chemically cross-linked by EDC/NHS endows collagen membrane with promising mechanical properties, it is not conducive to modulation of foreign body reaction (FBR) after implantation or guidance of osteogenesis. In our previous research, we have found that macrophages have a strong regulatory effect on tissue and bone regeneration during FBR, and EGCG modified membranes could adjust the recruitment and phenotypes of macrophages. Accordingly, we develop the EGCG-EDC/NHS membranes, prepared with physically immersion, while the surface morphology of the membrane was observed by SEM, the biological activity of collagen was determined by FTIR, the activity and adhesion of cell culture *in vitro*, angiogenesis and monocyte/macrophage recruitment after subcutaneous implantation, etc. are characterized. It could be concluded that EGCG-EDC/NHS collagen membrane is hopeful to be used in implant dentistry for it not only retains the advantages of the collagen membrane itself, but also improves cell viability, adhesion and vascularization tendency. However, the mechanism that lies in the regenerative advantages of such membrane needs further exploration, but it is certain that the differences in surface morphology can have a significant impact on the reaction between the host and the implant, not to mention macrophage in bone regeneration.

## 1. Introduction

Guided bone regeneration (GBR) technology takes advantage of barrier membrane to compartmentalize soft and hard tissue. It protects the hard tissue from an invasion of the relatively fast-growing soft tissue, thus creates sufficient space and provides stability for osteogenic cell migration in the meanwhile (Melcher, 1976). Currently, collagen, the main organic component of extracellular matrix (ECM) and bone, is the most common resorbable source for membrane fabrication. A meta-analysis revealed that the survival rate of implant with the application of collagen membrane were close to 100% both in simultaneous and subsequent implant placement (Wessing, Lettner, & Zechner, 2018). However, its high biocompatibility also becomes its fatal flaw, which limits the mechanical properties of the collagen membrane. Without the addition of crosslinking agents, the integrity of the membrane may not be able to maintain throughout the entire process of bone regeneration to provide sufficient support (Meyer, 2019). Although the exposure of membrane is slightly related to the addition of crosslinker, it is not statistically significant (30%) (Wessing et al., 2018). This observed degradation time is markedly longer than that of collagen membrane, which is reported to be completely resorbed 1 to 2 weeks after exposure [18, 34]. The prolonged degradation time of matrix barrier seems to provide prolonged protecting of the underlying graft supporting the bone regeneration process. During this healing process, all exposures did resolve within 6-7 weeks and no membrane had to be extracted. The ridge width gain in both groups was sufficient to allow for the successful placement of dental implants in all 14 subjects without any complication (Eskan, Girouard, Morton, & Greenwell, 2017). Therefore, to enhance mechanical properties with low cytotoxicity and low antigenicity, 1-ethyl-3-(3-dimethylaminopropylcarbodiimide hydrochloride (EDC) in the presence of N-hydroxy-succinimide (NHS) collagen membrane was selected as our target (Akhshabi, Biazar, Singh, Keshel, & Geetha, 2018; Bax et al., 2017). Apart from the degradation rate of membranes, immune environment of the implant site is another factor that determines the success or failure of the surgery (El-Jawhari, Jones, & Giannoudis, 2016). Especially in periodontitis, a chronic and degenerative inflammatory disease which affects approximately 10% of the world’s population, the inflammatory environment makes it harder to conduct tissue engineering with biomaterials, thus elevates the importance of immune modulation to rebuild balance (Kurashima & Kiyono, 2017). Epigallocatechin-3-gallate (EGCG) as one of the main polyphenols in tea could serve as a crosslinker for scaffolds (Honda et al., 2018), showing bioactive effects in various aspects, especially in dental restoration (Liao et al., 2020). Compared with pure collagen membrane, its anti-inflammatory (Lagha & Grenier, 2019; Wu, Choi, Kang, Kim, & Shin, 2017), anti-fibrosis (Wang, Yang, Yuan, Yang, & Zhao, 2018), pro-osteogenic (Lin et al., 2018), macrophage phenotypes regulatory (Chu et al., 2019) effects are all prompt to rescue the pro-long inflammation and help the establishment of ideal microenvironment to provide promising signal in regeneration (Chu et al., 2020). Considering its promising traits, EGCG was cross-linked to attach the commercial EDC/NHS collagen, and the modification of EGCG was adjusted at 0.064% w/v, where it is reported to possess appropriate mechanical properties, anti-inflammatory effects, and cell viability promotion in previous report (Chu et al., 2016). To ensure that the developed EGCG modified EDC/NHS collagen membrane meets the need of GBR, its physical, chemical and biological properties was characterized by means of investigation on surface morphology, FTIR spectra, DSC statistics, *in vitro* cell viability and integrin expression, and *in vivo* vascularization and monocyte/ macrophage recruitment. The aim of this study is to fabricate 0.064% EGCG EDC/NHS collagen membranes (**Fig. 1**) with the improvement of cell viability and adhesion, as well as revascularization.

**Fig. 1.**
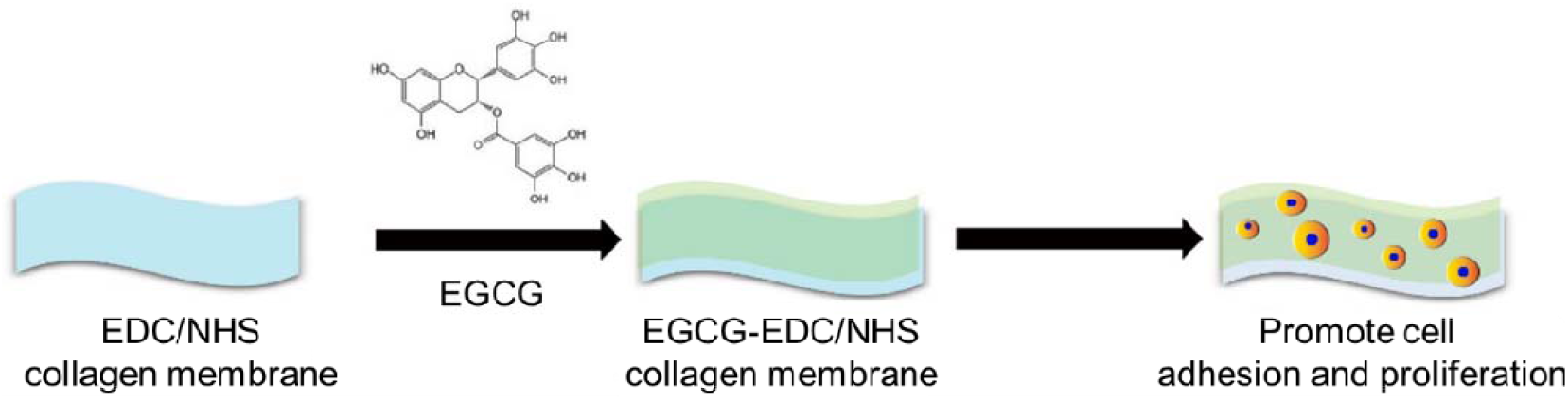
Schematic graph of loading 0.064% EGCG to EDC/NHS collagen membrane

## 2. Materials and Methods

### 2.1 Materials

Commercially available EDC/NHS collagen membranes (Dentium) in size of 10mm×20mm were purchased. EGCG was purchased from Jiang Xi Lv Kang Natural Products (Jiang Xi, China). All solvents and chemicals employed in the process were analytical grade and were used without further purification. The EGCG modified EDC/NHS collagen membrane was fabricated as follow: each EDC/NHS collagen membrane was immersed in 0.064% (w/v) EGCG solution at room temperature for 1 hr. After that, the EGCG-EDC/NHS collagen membranes were rinsed three times in deionized water and freeze dried overnight. The pure EDC/NHS collagen membranes were also processed under the same protocol except for immersion.

### 2.2 Surface morphology

The morphologies of EDC/NHS collagen membranes and EGCG modified collagen membranes were characterized by Scanning electron microscope (SEM, S-800, HITACHI, Tokyo, Japan) with an accelerating voltage of 25 kV. The discs were coated with an ultrathin layer (300 Å) of Au/Pt in an ion sputter (E1010, HITACHI, Tokyo, Japan) to achieve enough electrical conductivity.

### 2.3 Mechanical and chemical properties

#### a. Mechanical properties measurement

To investigate the ultimate stress (US), ultimate elongation (UE) and Young’s modulus (YM) of the membranes, each sample (10mm×20mm) was attached to an electronic universal testing machine (SHIMADZU, AG-IC 50 KN, Japan). Five samples were measured for each kind of membranes. Each sample was strained at a rate of 15mm/min at room temperature. When operating differential scanning calorimeter (DSC) measurement, samples in size of 10mm×20mm (∼6.71 mg) were encapsulated in aluminum pan, then heated from 25 to 500 °C at a rate of 10 °C/min under nitrogen atmosphere. Thermo-grams were obtained by the Netzsch Proteus analysis software, and thermal denaturation was recorded as the typical peak.

#### b. Chemical properties measurement

A Fourier transform infrared spectroscopy (FTIR) spectrophotometer (Spectrum One, PerkinElmer, Inc., Waltham) was employed to measure the FTIR spectra of the samples, the spectra of which were obtained at room temperature at the average of 32 scans in the range of 400 – 4000 cm−1.

### 2.4 Cell viability

The cell viability was examined using the mouse macrophage cell line Raw 264.7 obtained from the American Type Culture Collection (Rockville, MD, USA). Staining of Cell Counting Kit-8 (CCK-8, Dojindo Laboratories, Kumamoto, Japan) and Calcein AM/Hoechst were both conducted, but for different time periods. For CCK-8, membranes were processed into 10mm×10mm in the clean bench, then placed in 48-well plates and seeded with Raw 264.7 cell at a density of 10^4^ /well. 10% CCK-8 solution would be added to each well. Cells were then co-cultured for 10, 20, 40, 60, 120, 240, 360 min and 1 day, respectively, with RPMI (Gibco, Thermo, USA) supplemented with 10% FBS (Gibco, Thermo, USA). Then plates would be incubated at 37 °C for another 3.5 hr. After incubation, the OD value at 450 nm was measured to determine the cell viability through a micro-plate reader (Multiskan, Thermo, USA). As for staining of Calcein AM/Hoechst, cells were cultured for 5 days on the dishes then incubated with 1μM Calcein AM (diluted from a 1mM stock solution of CAM in dimethyl sulfoxide, Dojindo Laboratories, Kumamoto, Japan) and 100 μL RPMI for 30 mins in the incubator. The dishes were then washed three times by 1×PBS and stained with 100μL Hoechst 33258 working fluid (diluted from a 1mL stock solution in distilled water, KeyGen, China) for 10 mins in the incubator. At last, washed the dish for three times by 1×PBS and analyzed with an Inverted Ti-E microscope (Nikon).

### 2.5 Cell adhesion

SEM images were taken to preliminary estimate cell adhesion after culturing on EDC/NHS-Col and 0.064% EGCG-EDC/NHS-Col for 10, 20, 40 and 480 min, with processed as mentioned in 2.2. The expressions of integrin beta 1, 2, 3 in Raw 264.7 cells after culturing on blank wells, EDC/NHS-Col and 0.064% EGCG-EDC/NHS-Col for 10 mins were further evaluated by RNA isolation, cDNA synthesis, and RT-qPCR. Cells cultured on different surfaces were collected after incubating with 0.05% trypsin-EDTA (Gibco, Thermo, USA) for 5 mins and homogenized in 1-ml Trizol Reagent (Tianjian, Beijing, China). Total RNA was reverse-transcribed using mRNA Selective PCR kit (TaKaRa Bio-Clontech). Mouse integrin beta 1, 2 and 3 cDNA were amplified by real-time PCR using the SYBR Green PCR kit (Thermo Scientific, Braunschweig, Germany). The primer sequences used for the real-time PCR were listed in **Table 1**.

**Table 1.**
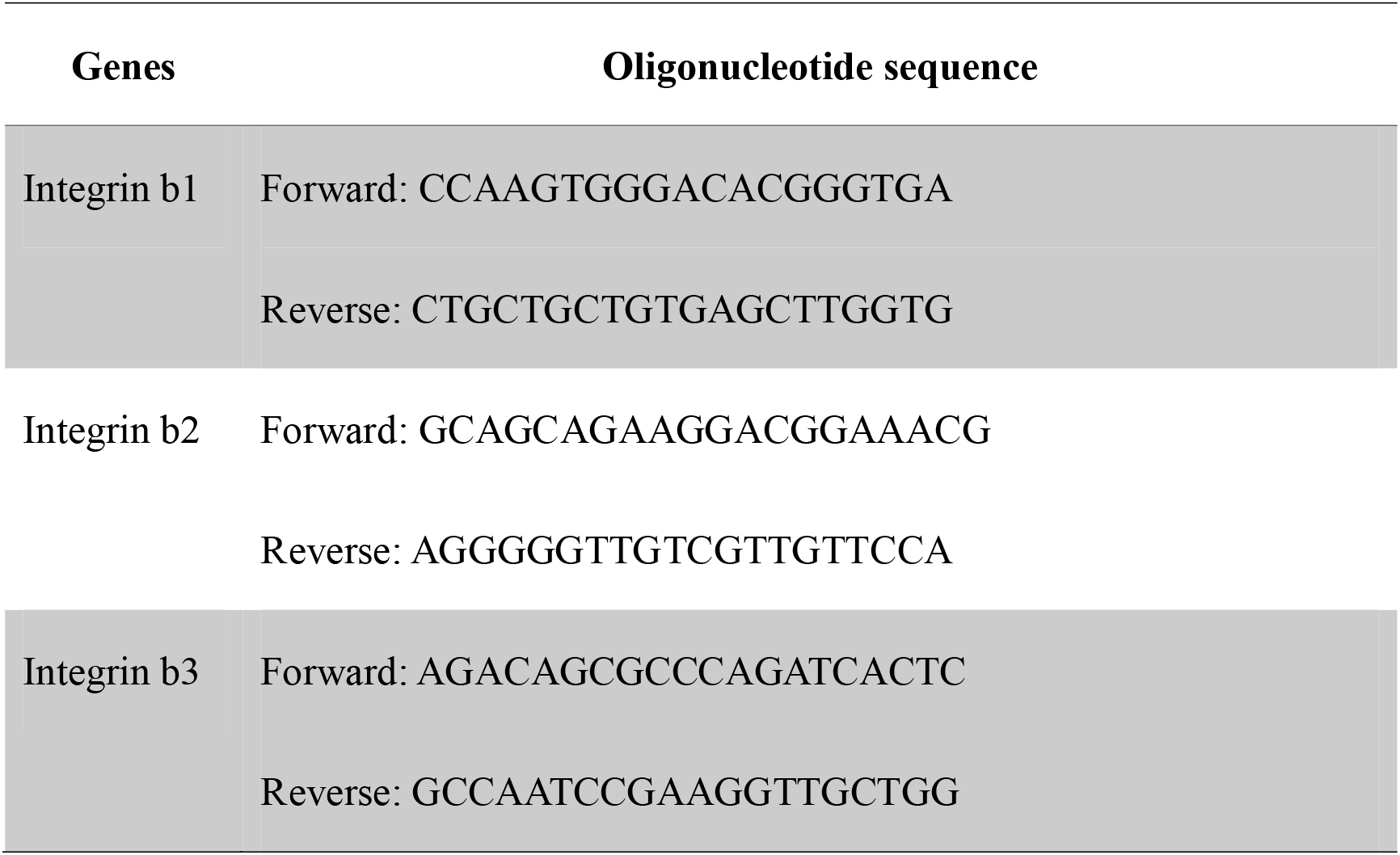
Nucleotide primers used for quantitative polymerase chain reaction.

### 2.6 Surgical Procedures

The protocol of the present experiment was approved by Institution Review Board of West China Hospital of Stomatology (No.WCHSIRB-D-2017-097). Male C57BL/6, 6∼7 weeks of age, were adaptively fed for 3 days after purchasing. After anaesthetization by chloral hydrate, the surgical area on the back was shaved and aseptically prepared. Three parallel sagittal incisions were made in the dorsal skin and subcutaneous pockets were prepared for membrane implantation. The two kinds of membranes were prepared beforehand into 3mm×3mm size in the clean bench. EDC/NHS collagen membranes and EGCG-EDC/NHS collagen membranes were implanted into the pockets respectively. The control group underwent a sham procedure. Afterwards, all the incisions were sutured. These mice were kept in a professional experimental animal room and fed with a standard laboratory diet. After recovering for 0 (12hr), 1, and 3 days, they were sacrificed by cervical dislocation. The membranes and the skins covering the materials were harvested together (equal areas around the suture were harvested in the control groups) and immediately fixed with 4% paraformaldehyde (Solarbio, Beijing, China) for at least 24hr.

### 2.7 H&E staining

The fixed samples were sliced into 3-μm thick sections after embedding in paraffin for the following H&E staining and immunofluorescence assay. For H&E staining, the sections were incubated at 65□ for 4hr for deparaffinization and processed as follow:

a. Dhydration with ethanol;
b. Stain with hematoxylin for 5 mins and differentiate in 1% hydrochloric acid alcohol for 2 s ;
c. Incubate in 0.2% ammonia water for 2 mins and stain with eosin for 1 min;
d. Gradual dehydration through 95% ethanol, then clear and mount the sections with neutral resin. The sections were observed with light microscopy (Olympus, Tokyo, Japan) and scanned with Digital Slide Scanning System (PRECICE, Beijing, China).

### 2.8 Promotion of revascularization

3 days after implantation, tissue was harvested and prepared as sections. Then, sections would be fixed with 2% paraformaldehyde PBS for 24 hours (pH=7.4) and later washed by PBS for 3 times, which followed by soaking in 1% bovine serum albumin (BSA) PBS that contained 1% Triton X-100 for 1 hour. Eventually, incubated the sections in 1% Tween 20 for 10 mins, and washed twice in PBS for 5 mins. The sections were then stained with nucleus marker Hoechst, endothelial cell marker CD31 and adventitial fibroblast marker α-SMA to visualized specific components in the fluorescent images.

### 2.9 Statistical Analysis

Data are presented as mean + standard deviation. Statistical computation was performed in GraphPad Prism 5.0 (GraphPad Software, San Diego, CA, USA) and statistical significance was analyzed using analysis of variance followed by Tukey’s multiple comparison tests. Semi-quantitative data of immunofluorescence staining and HE staining that did not follow a normal distribution were further analyzed with Mann-Whitney *U* test. Unless otherwise noted, *p*<0.05 was considered statistically significant.

## 3. Results

### 3.1 Surface morphology

As shown in representative SEM images (**Fig. 2**), the surface morphology of EDC/NHS collagen membranes exhibited a rough, disordered outlook, with fibers randomly distributed. After treating with 0.064% EGCG, there were no notable changes on the outlooks, while the arrangement of fibers appeared more ordered and intact. In addition, we also found that smaller fiber branches extend from the backbone when compared with those without EGCG treatment. These might be due to the loading of EGCG hydrogen bonds formation between EGCG molecules and collagen fibers.

**Fig. 2.**
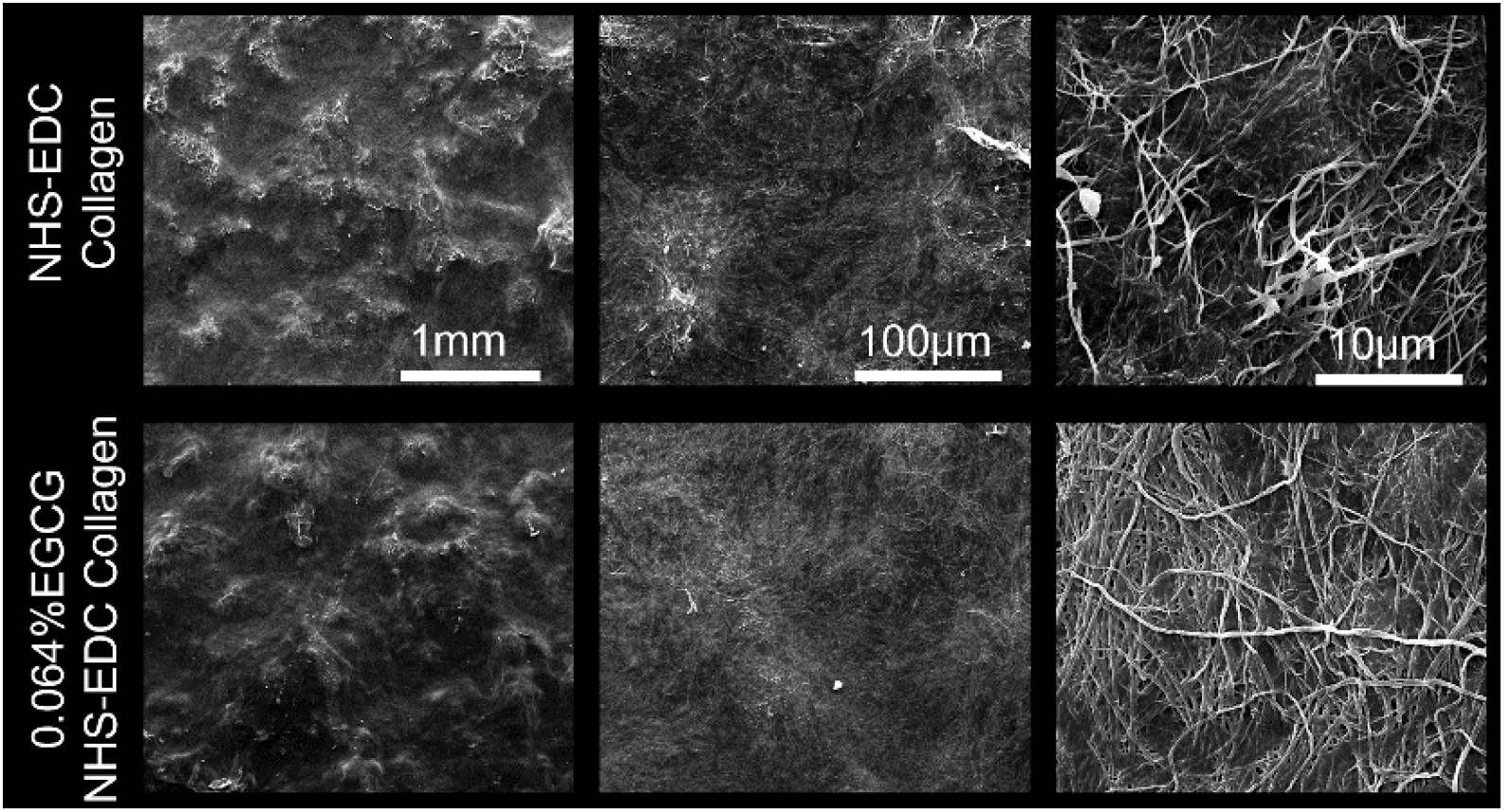
SEM images of EDC/NHS collagen membrane and 0.064% EGCG loaded EDC/NHS collagen membrane.

### 3.2 Mechanical and chemical properties

To examine the strength, stiffness and elasticity of the membranes, measurement of US, UE and YM were carried out and the corresponding volumes were written in **Table 2**. As US seemed relatively constant between the control and EGCG group, both UE and YM showed different trends. The modification of 0.064% EGCG strengthened the UE volume of collagen membranes, whereas weakened the YM volume (**Fig. 3a**). Also, there were no statistical difference between membranes with or without EGCG modification in result of DSC measurement (**Fig. 3b**). From results of FTIR spectra and DSC measurement, it could be concluded that chemical properties remain consistent before and after the treatment of 0.064% EGCG. The FTIR spectra indicated that no structure changes had happened to collagen fibers since the absorption peaked at 1235 cm^−1^ (tertiary structure) and 1450 cm^−1^ (pyrrolidine ring vibrations) of EDC/NHS collagen membrane, and 0.064% EGCG-EDC/NHS collagen membrane remained constant. Also, the ratio of 1235 cm^−1^/1450 cm^−1^ (**Table 3**) came out to be around 1.00, illustrating that the secondary structure of collagen triple helix remained its integrity in both membranes, which assured the great biological performance of collagen membranes (Figueiró, Góes, Moreira, & Sombra, 2004).Therefore, it could be inferred that EGCG did not affect the physicochemical property of collagen.

**Table 2.**
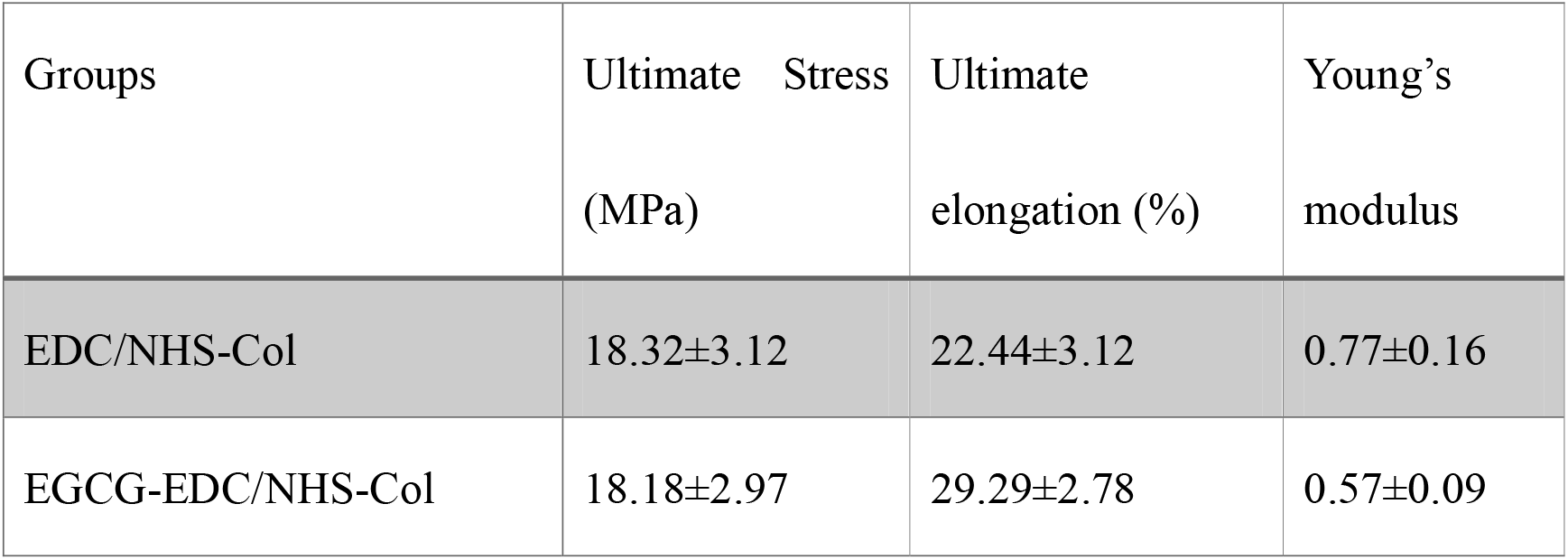
Mechanical properties of membranes.

| Groups | Ultimate Stress<br>(MPa) | Ultimate<br>elongation (%) | Young's<br>modulus |
| --- | --- | --- | --- |
| EDC/NHS-Col | 18.32±3.12 | 22.44±3.12 | 0.77±0.16 |
| EGCG-EDC/NHS-Col | 18.18±2.97 | 29.29±2.78 | 0.57±0.09 |

**Table 3.** FTIR ratio at the bands of 1235 cm^-1 and 1450 cm^-1

| Groups | 1235/1450 |
| --- | --- |
| NHS/EDC-Col | 1.001±0.002 |
| EGCG-NHS/EDC-Col | 0.989±0.003 |

**Fig. 3.**
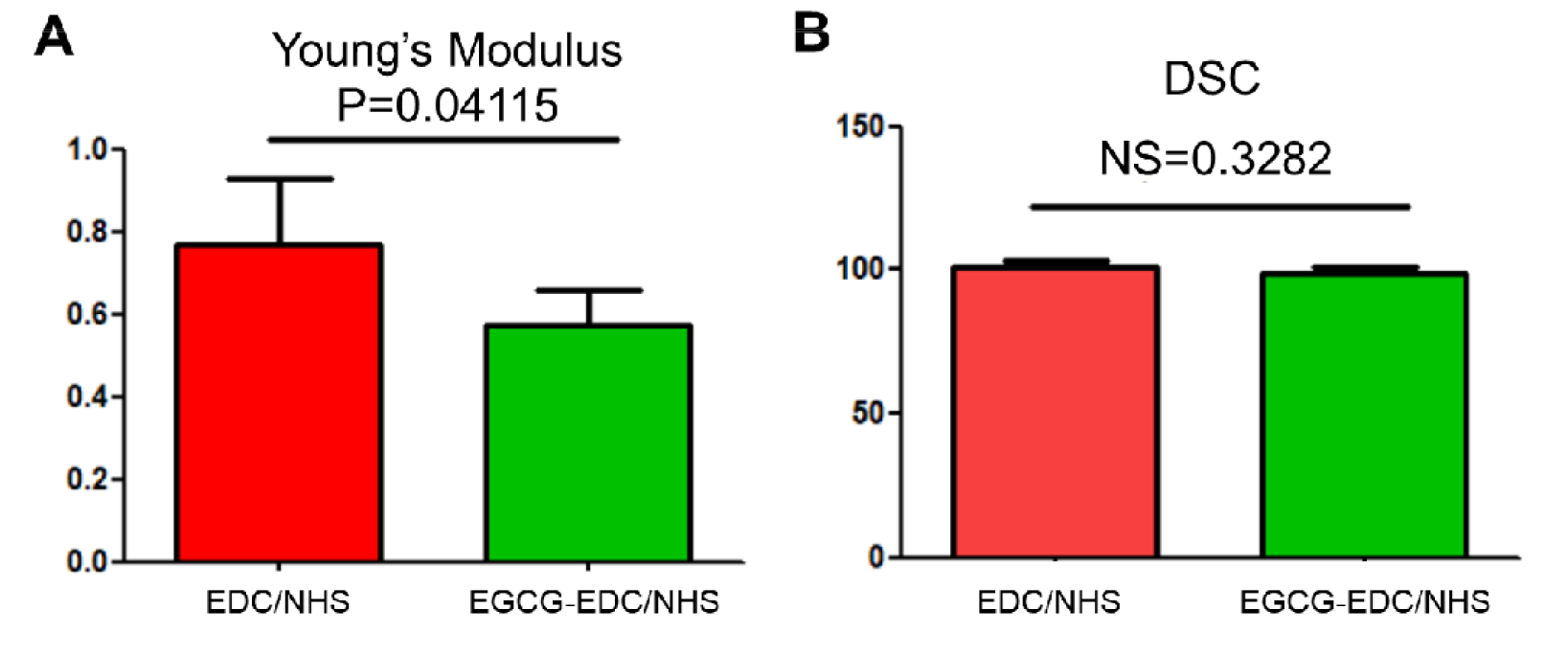
Young’s Modulus (a) and DSC statistics (b) of EDC/NHS collagen membrane and 0.064% EGCG-EDC/NHS collagen membrane. NS= no significance.

### 3.3 Cell Viability

The results of CCK-8 staining showed the cell viability of raw 264.7 cells under pure medium, 0.064% EGCG supplemented medium, EDC/NHS collagen membrane and 0.064% EGCG-EDC/NHS collagen membrane (**Fig. 4a**) from the first few minutes to 1 day. In the first hour, there was no significance between the groups. The improvement of cell viability by EGCG was raised after culturing for 2 hr and the following time points. In order to understand the effect of EGCG on cell viability after culturing for longer time period, Calcein AM/Hoechst staining was conducted on day 5 (**Fig. 4b**). The test results also showed the same trend with more live cells on EGCG-EDC/NHS-Col comparing with EDC/NHS-Col, indicating that the addition of EGCG was conducive to the performance of cell activities.

**Fig. 4.**
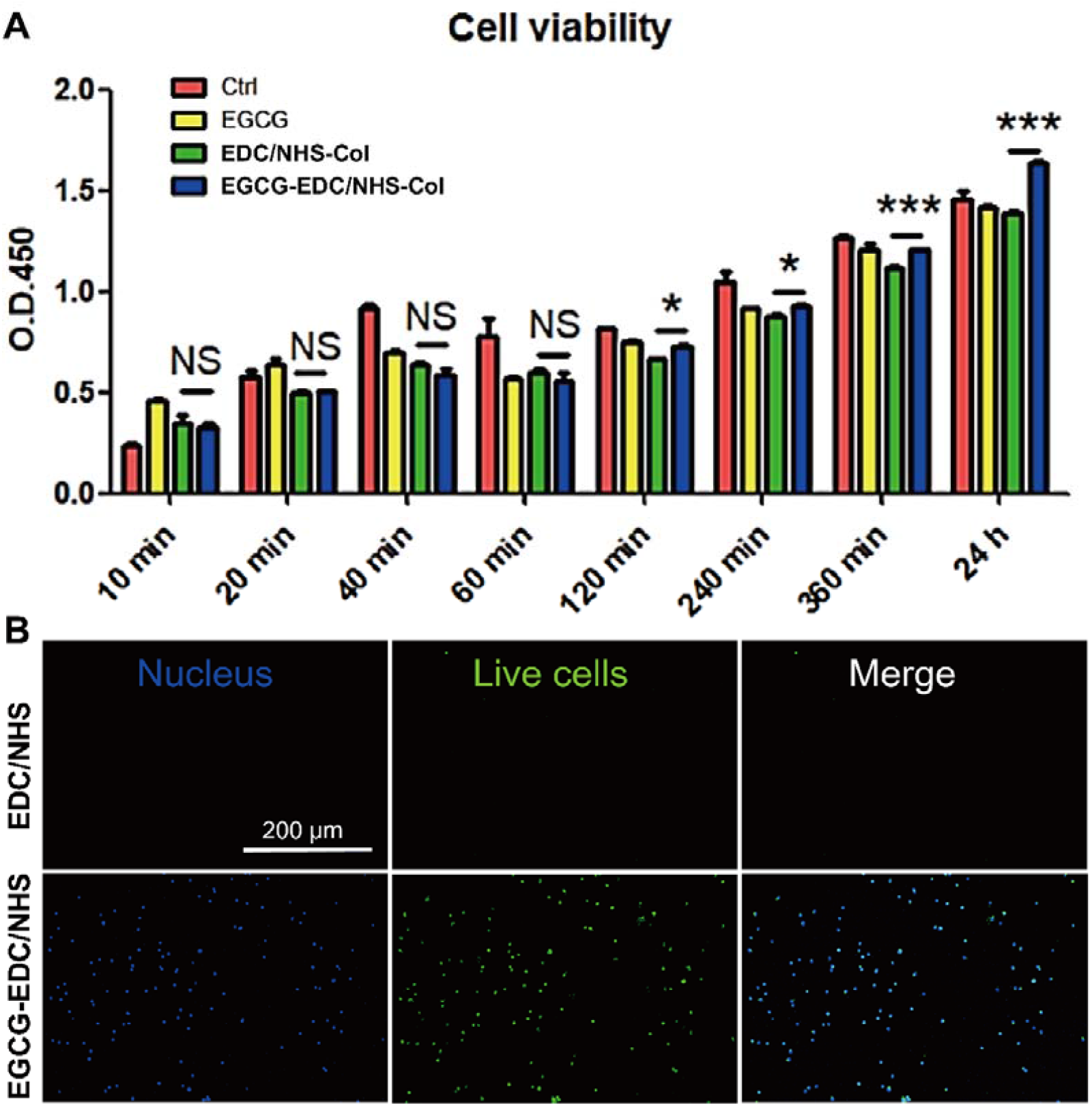
CCK-8 results (a) and Calcein AM/Hoechst staining (b) of Raw 264.7 cultured on different condition. Ctrl, standard medium; EGCG, 0.064% EGCG supplemented medium; EDC/NHS-Col, EDC/NHS collagen membrane; EGCG-EDC/NHS-Col, 0.064% EGCG-EDC/NHS collagen membrane. Green, live cells; blue, nucleus. NS= no significance, *P<0.05, **P<0.01, ***P<0.001, one-way ANOVA with Tukey’s multiple comparison test, n = 3 in each group.

### 3.4 Cell adhesion

To observe cell adhesion towards the membrane, SEM was used to generate the morphology of Raw 264.7 cells cultured on membranes in 8 hr (**Fig. 5a&b**). It elucidated that with the modification of EGCG, not only more cells adhered to the surface but also a better spreading condition of cells was detected. Also, more protrusions could be seen to seize the fibers even at very early incubation, compared to that without EGCG. Considering integrins, the transmembrane molecules, with large extracellular domains that are known to play a key role in adhesion between cells and ECM (Changede & Sheetz, 2017; Tseng et al., 2018), expression levels of integrin beta 1, 2 and 3 in Raw 264.7 cells were measured. The result showed that there existed a boost of integrin beta 2 in the EGCG-EDC/NHS group, whereas no differences appeared in the expression of integrin beta 1 and 3 with the treatment of 0.064% EGCG (**Fig. 5c**). These results suggested that 0.064% EGCG could enhance the adhesion properties of EDC/NHS collagen membrane at early phases.

**Fig. 5.**
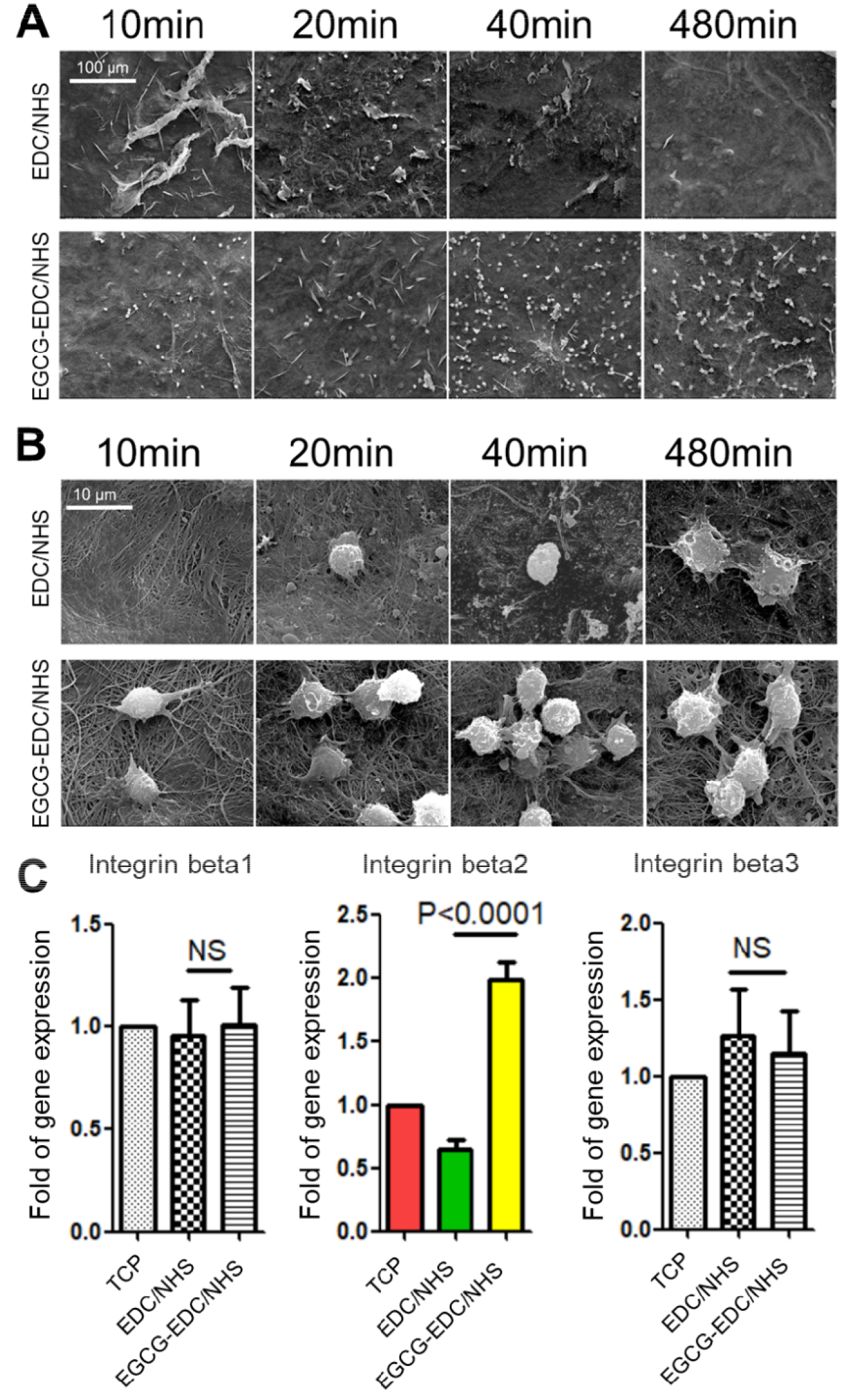
SEM images of Raw 264.7 cells cultured on EDC/NHS and EGCG treated EDC/NHS collagen membranes under low (a) and high (b) magnification, and the expression levels of integrin beta 1, 2 and 3, after culturing on tissue culture plate (TCP), EDC/NHS-Col and EGCG-EDC/NHS-Col for 10 mins (c), respectively. NS= no significance.

### 3.5 Promotion of revascularization

We identified the ability of 0.064% EGCG EDC/NHS-Col to promote revascularization by immunofluorescence staining of nuclei, vascular endothelial cell marker CD31 and adventitial fibroblast marker α-SMA (Manetti et al., 2017). As shown in **Fig. 6a**, more blood vessels were generated in EGCG-EDC/NHS-Col compared to EDC/NHS-Col, meanwhile, as shown in **Fig. 6d**, the results of semi-quantitative analysis of total blood vessel area showed that it was larger in 0.064% EGCG EDC/NHS-Col than that in the EDC/NHS-Col (p<0.01).

**Fig. 6.**
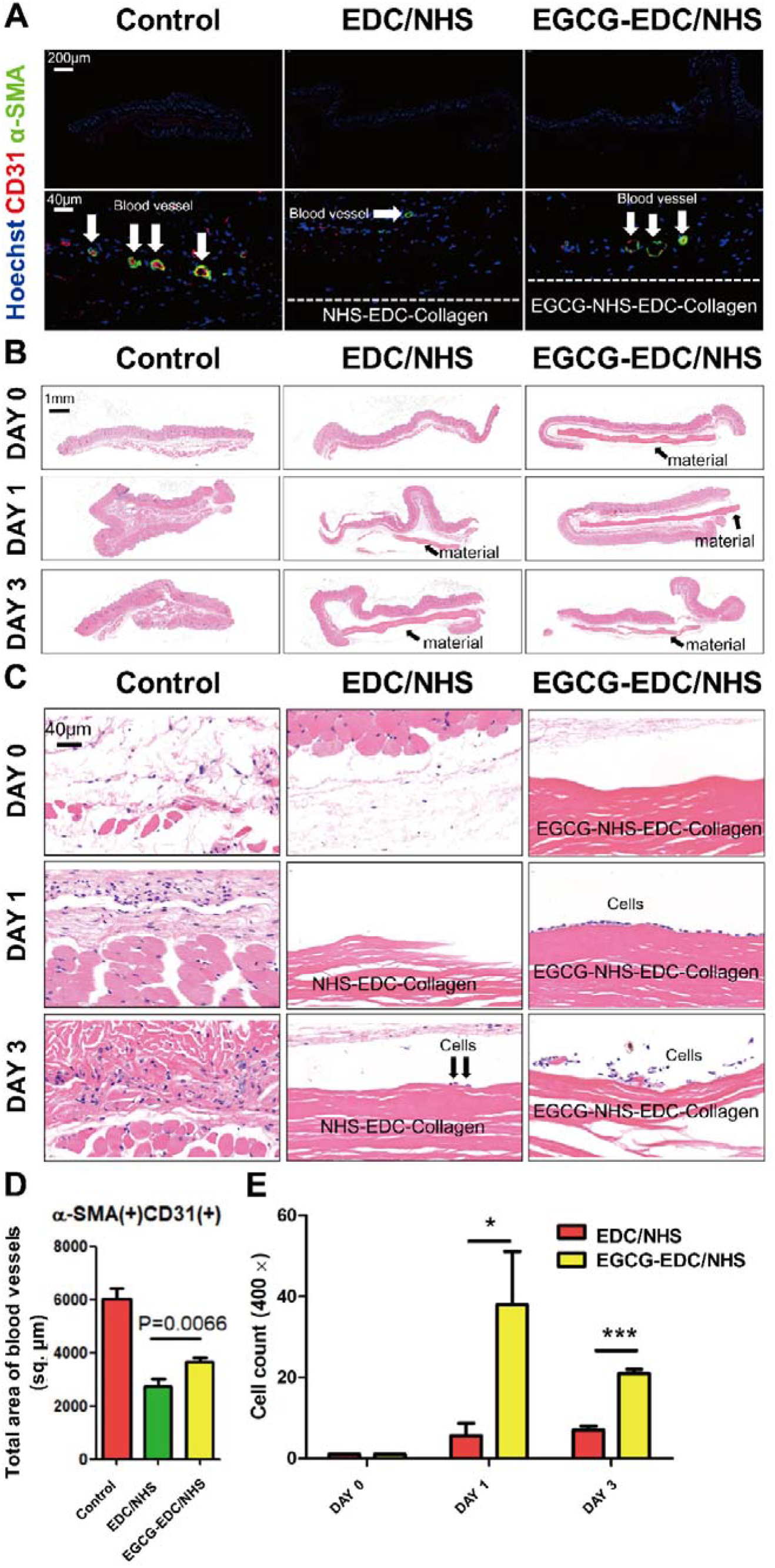
Results of subcutaneous implantation. Immunofluorescence staining of αSMA (green), CD31 (red) and Hoechst (blue) in subcutaneous implantation samples of EDC/NHS-Col and EGCG-EDC/NHS-Col, control group refers to normal skin (a); semi-quantitative analysis of total blood vessel area was calculated in each group (d). Recruitment of monocyte/macrophage after subcutaneous implantation of EDC/NHS collagen membrane and EGCG-EDC/NHS collagen membrane, control group underwent a sham procedure. HE staining of subcutaneous tissue (b&c) and cell counting of monocyte/macrophage (e) in NHS/EDC-Col and EGCG-EDC/NHS-Col were conducted on day 0, 1, and 3 post implantations. Arrows indicate material (b) and monocyte/macrophage (c). Statistical significance was analyzed using analysis of variance followed by Tukey’s multiple comparison tests and Mann-Whitney *U* test (*n* = 5). Data are presented as mean + standard deviation. *P<0.05, ***P<0.001.

### 3.6 Recruitment of monocyte/macrophage

12 hours after the implantation, there were no visible cells recruited at surgical sites, but with another 12 hours, we could see that around the membrane, a certain number of immune cells were gathered, which was similar on 3-days post implantation (**Fig. 6c&d**). In addition, semi-quantitative cell counting revealed that in 1- and 3-days post surgeries, 0.064% EGCG-EDC/NHS-Col recruited more monocytes/macrophages, while EDC/NHS-Col was about one-third of the number of the experimental group (**Fig. 6e**).

## 4. Discussion

To reconstruct alveolar bone deficiency, the application of biomaterials and certain surgery are on account of its morphology and the severity of horizontal/ vertical bone loss. When performing GBR, bone defect less than 6 mm is a prerequisite for desirable outcome, whereas the complex case that is combined with vertical and horizontal bone defect ≤ 4mm, it is recommended to use crosslinked absorbable membrane or non-absorbable membrane with bone filler to create appropriate repair conditions (Tolstunov, Hamrick, Broumand, Shilo, & Rachmiel, 2019). In addition, crosslinked membranes are more osteoinductive, with more bone marrow-multipotent stromal cells attachment to the crosslinked membranes, better ALP activity, and calcium deposition were observed compared with the non-crosslinked one (El-Jawhari, Moisley, Jones, & Giannoudis, 2019). According to previous studies, the use of EDC as crosslinker could not only prolong the integrity of the membrane, but also avoid severe FBR, lack of vascularization in the early stage of healing and poor integration, which are the fatal flaws of traditional cross-linking agents (Park et al., 2015). The addition of NHS would improve EDC-mediated crosslinking, stabilize active intermediates, and reduce side products for subsequent reactions (Grabarek & Gergely, 1990). However, the present EDC/NHS crosslinked collagen membrane lacks the ability to regulate immune responses and induce osteoblast-related cell behavior. Regarding cell behavior, EGCG has shown the ability to complement these two aspects in our previous research as mentioned in the introduction. Thus, we attempted to modify to EDC/NHS crosslinked collagen membrane with 0.064% EGCG. After characterizing the material, results show that the backbone of collagen remains intact and the strength of membranes is barely altered, whereas the stiffness is moderately enhanced, though elasticity is slightly weakened. These not only ensure that the membrane will not be too supple to tightly cover the substitutes, supporting bone formation, but also verify that the chemical structure related to the biological activity of collagen has not been destroyed, though the definite clinical effect needs verification. Over the past decade, the ability of biomaterials to promote tissue regeneration by regulating immune cells in advance has been confirmed, the composition, physical and chemical properties, and surface morphology of which are the critical factors that affect the foreign body reaction (FBR) after implantation. As one of the dominant immune cells in the FBR, macrophages can acutely polarize into different phenotypes according to the microenvironment created by the implant and direct the accumulated cells’ behaviors that strongly involve in tissue reconstruction (Chu, Deng, Sun, Qu, & Man, 2017; Xie et al., 2020). The arrangement and diameter of the fibers can significantly affect the behavior of macrophages. Anisotropic membranes with thicker fibers have an increased tendency to oxidative degradation, compared with the thinner and isotropic one (Wissing et al., 2019), more macrophages adherence on the align compared with the random and the smaller fibers also have been proved with better biocompatibility as thinner fibrous capsule and abundant volume of blood vessel formation (Saino et al., 2011). In this article, we found that the surface morphology of EGCG-EDC/NHS-Col has been altered with smaller fiber branches extended from the backbone and the arrangement became more coherent, which may account for the cell viability and adherence. Under the electron microscope, the viability of RAW 264.7 on EGCG-EDC/NHS-Col is significantly higher than any other groups at all detection points during 2-24 hours after implantation. Not only did more macrophages adhere to membrane with the treatment of EGCG, but the adhered macrophages were activated at an early stage (20min) with many protrusions, and which is also confirmed by PCR detection. Also, more monocyte/macrophage recruitment could be seen on EGCG-EDC/NHS-Col in the *in vivo* subcutaneous implantation. As researchers have found that the onset of neovascularization greatly depends on macrophage and its coherent phenotypic switch (Spiller et al., 2014; Spiller, Freytes, & Vunjak-Novakovic, 2015), we hypothesize the promising angiogenesis is result from the recruitment of macrophages which cannot be excluded considering the microstructure of EGCG-EDC/NHS-Col. Of note, in our previous study, the modification of EGCG was highly competent at promoting vascularization that involved the secretion of M2-related chemokines (Chu et al., 2018). The formation of blood vessel could offer nutrient, stem cells and oxygen supply and waste discharge, vastly improving the regenerative response. Therefore, our efforts are directed at macrophages and the outcome of vascularization as primary study towards the developed EGCG modified EDC/NHS collagen membrane. And our results showed that the modification of EGCG is beneficial for cell viability, adhesion and vessel formation in both *in vitro* RAW 264.7 culture experiment and in vivo subcutaneous implantation. The great biocompatibility of EGCG modified EDC/NHS collagen guided bone regeneration membrane is stated without compromising the advantages of collagen itself. Furthermore, our results demonstrated the modified membranes could significantly affect the attachment of macrophages and enhance its viability both *in vivo* and *in vitro*, which might be related to the formation of vessels. However, the phenotypes of macrophages and mechanisms of angiogenesis need further investigation. In short, we develop a biomaterial that can promote the early recruitment of macrophages and the formation of blood vessels, possessing great potential for bone regeneration in the field of implant dentistry.

## 5. Conclusion

In conclusion, with the modification of EGCG, EDC/NHS collagen guided bone regeneration membrane can better promote cell adhesion and improve cell viability, which is confirmed in our experiments that living cells attach to it and the expression of adhesion-related integrin is indeed increased. In addition, there was a statistically significant difference in angiogenesis after membrane implantation. The benefits mentioned above do not exclude the presence of smaller collagen fibers, but this different microstructure does not damage the chemical structure of the collagen itself, though the specific effect needs further exploration.

## Reference

Akhshabi, S., Biazar, E., Singh, V., Keshel, S. H., & Geetha, N. (2018). The effect of the carbodiimide cross-linker on the structural and biocompatibility properties of collagen-chondroitin sulfate electrospun mat. International journal of nanomedicine, 13, 4405–4416.

Bax, D. V., Davidenko, N., Gullberg, D., Hamaia, S. W., Farndale, R. W., Best, S. M., & Cameron, R. E. (2017). Fundamental insight into the effect of carbodiimide crosslinking on cellular recognition of collagen-based scaffolds. Acta biomaterialia, 49, 218–234.

Changede, R., & Sheetz, M. (2017). Integrin and cadherin clusters: A robust way to organize adhesions for cell mechanics. BioEssays : news and reviews in molecular, cellular and developmental biology, 39(1), 1–12.

Chu, C., Deng, J., Sun, X., Qu, Y., & Man, Y. (2017). Collagen Membrane and Immune Response in Guided Bone Regeneration: Recent Progress and Perspectives. Tissue engineering. Part B, Reviews, 23(5), 421–435.

Chu, C., Deng, J., Xiang, L., Wu, Y., Wei, X., Qu, Y., & Man, Y. (2016). Evaluation of epigallocatechin-3-gallate (EGCG) cross-linked collagen membranes and concerns on osteoblasts. Materials science & engineering. C, Materials for biological applications, 67, 386–394.

Chu, C., Liu, L., Rung, S., Wang, Y., Ma, Y., Hu, C., … Qu, Y. (2020). Modulation of foreign body reaction and macrophage phenotypes concerning microenvironment. Journal of biomedical materials research. Part A, 108(1), 127–135.

Chu, C., Liu, L., Wang, Y., Wei, S., Wang, Y., Man, Y., & Qu, Y. (2018). Macrophage phenotype in the epigallocatechin-3-gallate (EGCG)-modified collagen determines foreign body reaction. Journal of Tissue Engineering and Regenerative Medicine, 12(6), 1499–1507.

Chu, C., Wang, Y., Wang, Y., Yang, R., Liu, L., Rung, S., … Qu, Y. (2019). Evaluation of epigallocatechin-3-gallate (EGCG) modified collagen in guided bone regeneration (GBR) surgery and modulation of macrophage phenotype. Materials science & engineering. C, Materials for biological applications, 99, 73–82.

El-Jawhari, J. J., Jones, E., & Giannoudis, P. V. (2016). The roles of immune cells in bone healing; what we know, do not know and future perspectives. Injury-international Journal of the Care of the Injured, 47(11), 2399–2406.

El-Jawhari, J. J., Moisley, K., Jones, E., & Giannoudis, P. V. (2019). A crosslinked collagen membrane versus a non-crosslinked bilayer collagen membrane for supporting osteogenic functions of human bone marrow-multipotent stromal cells. European cells & materials, 37, 292–309.

Eskan, M. A., Girouard, M. E., Morton, D., & Greenwell, H. (2017). The effect of membrane exposure on lateral ridge augmentation: a case-controlled study. International journal of implant dentistry, 3(1), 26.

Figueiró, S. D., Góes, J. C., Moreira, R. A., & Sombra, A. S. B. (2004). On the physico-chemical and dielectric properties of glutaraldehyde crosslinked galactomannan–collagen films. Carbohydrate Polymers, 56(3), 313–320.

Grabarek, Z., & Gergely, J. (1990). Zero-length crosslinking procedure with the use of active esters. Analytical biochemistry, 185(1), 131–135.

Honda, Y., Takeda, Y., Li, P., Huang, A., Sasayama, S., Hara, E., … Tanaka, T. (2018). Epigallocatechin Gallate-Modified Gelatin Sponges Treated by Vacuum Heating as a Novel Scaffold for Bone Tissue Engineering. Molecules, 23(4), 876.

Kurashima, Y., & Kiyono, H. (2017). Mucosal Ecological Network of Epithelium and Immune Cells for Gut Homeostasis and Tissue Healing. Annual review of immunology, 35, 119–147.

Lagha, A. B., & Grenier, D. (2019). Tea polyphenols protect gingival keratinocytes against TNF-α-induced tight junction barrier dysfunction and attenuate the inflammatory response of monocytes/macrophages. Cytokine, 115, 64–75.

Liao, S., Tang, Y., Chu, C., Lu, W., Baligen, B., Man, Y., & Qu, Y. (2020). Application of green tea extracts epigallocatechin-3-gallate in dental materials: Recent progress and perspectives. Journal of biomedical materials research. Part A.

Lin, S. Y., Kang, L., Wang, C. Z., Huang, H. H., Cheng, T. L., Huang, H. T., … Chen, C. H. (2018). (-)-Epigallocatechin-3-Gallate (EGCG) Enhances Osteogenic Differentiation of Human Bone Marrow Mesenchymal Stem Cells. Molecules, 23(12), 3221.

Manetti, M., Romano, E., Rosa, I., Guiducci, S., Bellando-Randone, S., De Paulis, A., … Matucci-Cerinic, M. (2017). Endothelial-to-mesenchymal transition contributes to endothelial dysfunction and dermal fibrosis in systemic sclerosis. Annals of the rheumatic diseases, 76(5), 924–934.

Melcher, A. H. (1976). On the repair potential of periodontal tissues. Journal of periodontology, 47(5), 256–260.

Meyer, M. (2019). Processing of collagen based biomaterials and the resulting materials properties. Biomedical engineering online, 18(1), 24.

Park, J. Y., Jung, I. H., Kim, Y. K., Lim, H. C., Lee, J. S., Jung, U. W., & Choi, S. H. (2015). Guided bone regeneration using 1-ethyl-3-(3-dimethylaminopropyl) carbodiimide (EDC)-cross-linked type-I collagen membrane with biphasic calcium phosphate at rabbit calvarial defects. Biomaterials research, 19, 15.

Saino, E., Focarete, M. L., Gualandi, C., Emanuele, E., Cornaglia, A. I., Imbriani, M., & Visai, L. (2011). Effect of electrospun fiber diameter and alignment on macrophage activation and secretion of proinflammatory cytokines and chemokines. Biomacromolecules, 12(5), 1900–1911.

Spiller, K. L., Anfang, R. R., Spiller, K. J., Ng, J., Nakazawa, K. R., Daulton, J. W., & Vunjak-Novakovic, G. (2014). The role of macrophage phenotype in vascularization of tissue engineering scaffolds. Biomaterials, 35(15), 4477–4488.

Spiller, K. L., Freytes, D. O., & Vunjak-Novakovic, G. (2015). Macrophages modulate engineered human tissues for enhanced vascularization and healing. Annals of biomedical engineering, 43(3), 616–627.

Tolstunov, L., Hamrick, J. F. E., Broumand, V., Shilo, D., & Rachmiel, A. (2019). Bone Augmentation Techniques for Horizontal and Vertical Alveolar Ridge Deficiency in Oral Implantology. Oral and maxillofacial surgery clinics of North America, 31(2), 163–191.

Tseng, H. Y., Samarelli, A. V., Kammerer, P., Scholze, S., Ziegler, T., Immler, R., … Böttcher, R. T. (2018). LCP1 preferentially binds clasped αMβ2 integrin and attenuates leukocyte adhesion under flow. Journal of cell science, 131(22), jcs218214.

Wang, L., Yang, G., Yuan, L., Yang, Y., & Zhao, H. (2018). Green Tea Catechins Effectively Altered Hepatic Fibrogenesis in Rats by Inhibiting ERK and Smad1/2 Phosphorylation. Journal of agricultural and food chemistry, 67(19), 5437–5445

Wessing, B., Lettner, S., & Zechner, W. (2018). Guided Bone Regeneration with Collagen Membranes and Particulate Graft Materials: A Systematic Review and Meta-Analysis. The International journal of oral & maxillofacial implants, 33(1), 87–100.

Wissing, T. B., Bonito, V., van Haaften, E. E., van Doeselaar, M., Brugmans, M., Janssen, H. M., … Smits, A. (2019). Macrophage-Driven Biomaterial Degradation Depends on Scaffold Microarchitecture. Frontiers in bioengineering and biotechnology, 7, 87.

Wu, Y. R., Choi, H. J., Kang, Y. G., Kim, J. K., & Shin, J. W. (2017). In vitro study on anti-inflammatory effects of epigallocatechin-3-gallate-loaded nano- and microscale particles. International journal of nanomedicine, 12, 7007–7013.

Xie, Y., Hu, C., Feng, Y., Li, D., Ai, T., Huang, Y., … Tan, J. (2020). Osteoimmunomodulatory effects of biomaterial modification strategies on macrophage polarization and bone regeneration. Regenerative biomaterials, 7(3), 233–245.

